# Refining the role of *de novo* protein truncating variants in neurodevelopmental disorders using population reference samples

**DOI:** 10.1101/052886

**Authors:** Jack A. Kosmicki, Kaitlin E. Samocha, Daniel P. Howrigan, Stephan J. Sanders, Kamil Slowikowski, Monkol Lek, Konrad J. Karczewski, David J. Cutler, Bernie Devlin, Kathryn Roeder, Joseph D. Buxbaum, Benjamin M. Neale, Daniel G. MacArthur, Dennis P. Wall, Elise B. Robinson, Mark J. Daly

## Abstract

Recent research has uncovered an important role for *de novo* variation in neurodevelopmental disorders. Using aggregated data from 9246 families with autism spectrum disorder, intellectual disability, or developmental delay, we show ~1/3 of *de novo* variants are independently observed as standing variation in the Exome Aggregation Consortium’s cohort of 60,706 adults, and these *de novo* variants do not contribute to neurodevelopmental risk. We further use a loss-of-function (LoF)-intolerance metric, pLI, to identify a subset of LoF-intolerant genes that contain the observed signal of associated *de novo* protein truncating variants (PTVs) in neurodevelopmental disorders. LoF-intolerant genes also carry a modest excess of inherited PTVs; though the strongest *de novo* impacted genes contribute little to this, suggesting the excess of inherited risk resides lower-penetrant genes. These findings illustrate the importance of population-based reference cohorts for the interpretation of candidate pathogenic variants, even for analyses of complex diseases and *de novo* variation.

## Introduction

Autism spectrum disorders (ASDs) are a phenotypically heterogeneous group of heritable disorders that affect ~1 in 68 individuals in the United States^1^. While estimates of the common variant (heritable) contribution toward ASD liability are upwards of 50%^2-4^, few specific risk variants have been identified, in part because ASD GWAS sample sizes to date remain limited. Conversely, the field made substantial progress understanding the genetic etiology of ASD via analysis of *de novo* (newly arising) variation using exome sequencing of parent-offspring trios^5-10^. Severe intellectual disability and developmental delay (ID/DD) are considerably less heritable than ASDs^11^ (though frequently comorbid) and have demonstrated a stronger contribution from *de novo* frameshift, splice acceptor, splice donor, and nonsense variants (collectively termed protein truncating variants [PTVs])^12-14^. Furthermore, ASD cases with comorbid ID show a significantly higher rate of *de novo* PTVs than those with normal or above average IQ^6,9,15-17^, while higher IQ cases have a stronger family history of neuropsychiatric disease^15^, suggesting a greater heritable contribution.

*De novo* variants comprise a unique component of the genetic architecture of human disease since, having not yet passed through a single generation, any heterozygous variants with complete or near-complete elimination of reproductive fitness must reside almost exclusively in this category. Despite prior evidence of mutational recurrence^18^ (i.e., the same mutation occurring *de novo* in multiple individuals), most studies implicitly assumed each *de novo* variant was novel, in line with Kimura’s infinite sites model^19^, and thereafter analyzed *de novo* variants genome-wide without respect to their allele frequency in the population (**Supplementary Note**). However, the mutation rate is not uniform across the genome, with some regions and sites experiencing higher mutation rates than others (e.g., CpG sites^20^). A classic example comes from achondroplasia, in which the same G-to-C and G-to-T variant at a CpG site was observed *de novo* in 150 and 3 families, respectively^18^. As such, it should not be surprising to observe a *de novo* variant at a given site and also observe the same variant (defined as one with the same chromosome, position, reference, and alternate allele) present as standing variation in the population.

Given the strong selective pressure on neurodevelopmental disorders^21-23^, we expect most highly deleterious (high-risk conferring) *de novo* PTVs will linger in the population for at most a few generations. Thus, the collective frequency of such variants in the population will approximate their mutation rate. Individual PTVs tolerated to be seen in relatively healthy adults, and more generally PTVs in genes that tolerate the survival of such variants in the population, may be less likely to contribute significant risk to such phenotypes, and are therefore permitted by natural selection to reach allele frequencies orders of magnitude larger than those of highly deleterious variants. Given the current size of the human population (~7 billion), and the expectation of one *de novo* variant per exome (1 in ~30 million bases), every non-embryonic lethal coding mutation is likely present as a *de novo* variant at least once in the human population. This reasoning, along with the availability of large exome sequencing reference databases, motivated our interest in searching for variants observed *de novo* in trio sequencing studies that are also present as standing variation in the human population, indicating a recurrent mutation. We will herein refer to these *de novo* variants that are also observed as standing variation in the population as class 2 *de novo* variants, with the remaining *de novo* variants referred to as class 1 *de novo* variants (i.e., observed only *de novo;* **Fig. 1**).

**Figure 1.**
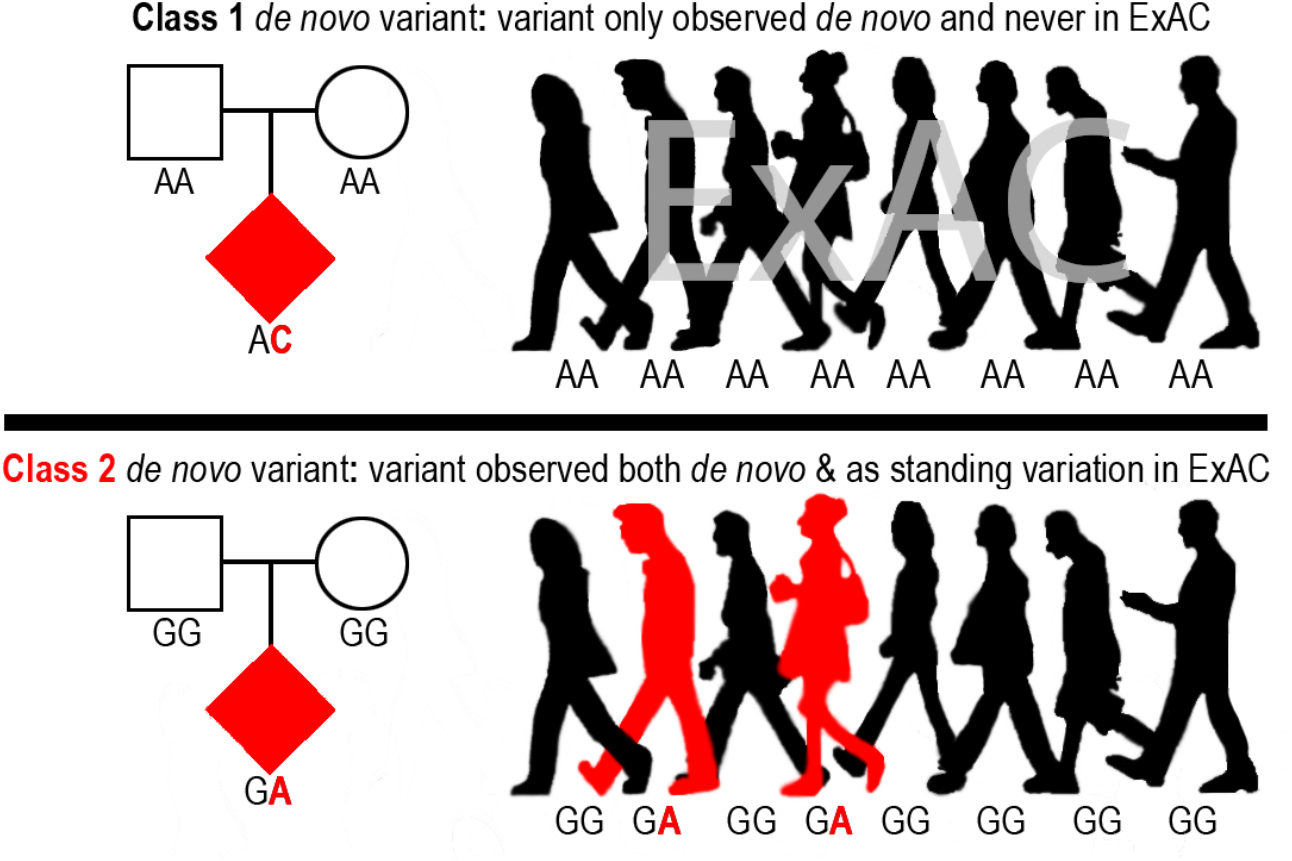
Illustration of class 1 and class 2 *de novo* variants with the genotypes of each variant for 8 of the 60,706 individuals in ExAC.

With the release of the Exome Aggregation Consortium’s (ExAC) dataset of 60,706 adult individuals without severe developmental abnormalities^24^, we can now empirically investigate the rate and relative pathogenicity of recurrent mutations. While there have been many studies examining *de novo* variation in human disease, the success in ASD and ID/DD, coupled with the large sample sizes published to date, led us to focus our evaluation on these phenotypes.

## Results

### Class 2 *de novo* variation

We first asked how many of the 10,093 variants observed *de novo* in ID/DD cases^13^, ASD cases, and unaffected ASD siblings are also observed as standing variation in the 60,706 reference exomes from ExAC^24^ **(Fig. 1;** Online Methods). We found that 1854 ASD (31.66%), 841 unaffected ASD sibling (33.05%), and 410 ID/DD (24.23%) *de novo* variants were observed as standing variation in one or more ExAC individuals (class 2 *de novo* variants) **(Fig. 2A; Supplementary Tables 1–3).** When we removed the 15,330 exomes originating from psychiatric cohorts (many of which are controls), the rate of class 2 *de novo* variation drops to 28.47% (±1.03%, 95% CI), a rate statistically indistinguishable from the expected recurrence rate of 28.13% (±0.42%, 95% CI; two-sided binomial test; *P*=0.45; **Fig. 2B; Supplementary Figs. 1 & 2; Supplementary Table 4;** Online Methods). We found similar rates of class 2 *de novo* variants in published trio studies of schizophrenia^25^ and congenital heart disease^26,27^ **(Supplementary Tables 5 & 6).** While the presence of class 2 *de novo* variants is not a novel observation^18,25^, the rate is approximately three times larger than previous estimates^25^ owing to significantly larger reference datasets **(Fig. 2B; Supplementary Fig. 2).**

We first sought confirmation that the observed recurrence rate – the proportion of variants observed both *de novo* and as standing variation in the population – was technically sound and not the result of some undetected contamination or missed heterozygote calls in parents (i.e., false *de novo* variants). Five secondary analyses strongly support the technical validity of this work. 1) In line with previous publications^25^, class 2 *de novo* variants, regardless of their functional impact, are enriched at CpG sites as compared to class 1 *de novo* variants (*P* < 1x10^−20^; Fisher’s exact test; **Supplementary Table 7)**. 2) As the exomes used to call *de novo* variants in De Rubeis et al. (2014) were used in the joint calling of ExAC, and many were sequenced at the same center as the majority of ExAC samples, it is possible that false class 2 *de novo* variants could be elevated in this dataset via contamination or joint calling artifacts. However, we observe no difference in the rate of class 2 *de novo* variation between De Rubeis et al. (2014) and Iossifov, O’Roak, Sanders, Ronemus et al. (2014) (*P*=0.10; Fisher’s exact test; **Supplementary Table 8**). 3) The frequency distribution of class 2 *de novo* variants should be skewed dramatically upward towards common variation if contamination or missed parental heterozygotes were contributing; however, the ExAC frequency of class 2 *de novo* variants at CpGs were indistinguishable from all such variants in ExAC compared to variants drawn at random with the same annotation and CpG content (*P*=0.14; Wilcoxon rank sum test; **Fig. 2C**). 4) In fact, a subset of synonymous variants experimentally validated in the ASD studies showed nearly the same recurrence rate as the overall set, most definitively establishing that these mutations indeed arose independently (*P*=0.60; Fisher’s exact test). 5) Lastly, it is well documented that mutation rate increases with paternal age^28-31^, thus rates of both class 1 and class 2 *de novo* variants should show association with paternal age if both classes were genuine *de novo* variants. Indeed, for the 1861 unaffected ASD siblings with available paternal age information, rates of both class 1 and class 2 *de novo* variants are associated with increasing paternal age (class 1: β=0.002, *P*=4.11x10^−9^; class 2: β=0.0009, *P*=1.06x10^−5^; linear regression).

**Figure 2.**
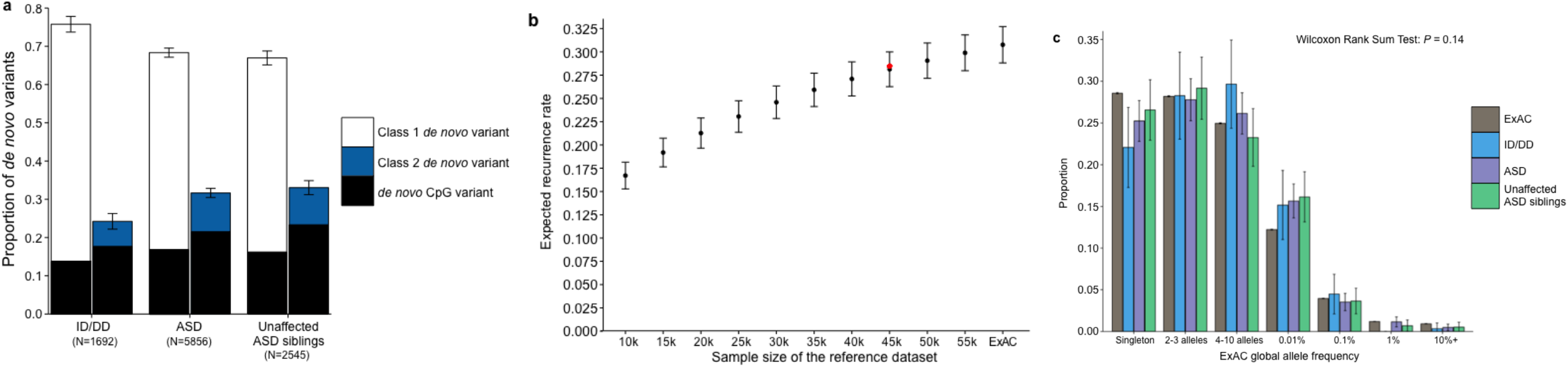
Properties of class 2 *de novo* variants. **(a)** The proportion of *de novo* variants across each cohort split between class 1 (left) and class 2 (right) with CpG variants marked in black. Class 2 *de novo* variants are strongly enriched for CpG variants (*P* < 10^−20^). The corresponding figure using the non-psychiatric version of ExAC can be found in **Supplementary Figure 2**. (**b**) Expected recurrence rate (rate of class 2 *de novo* variants across ID/DD, ASD, and unaffected ASD siblings) given the sample size of the reference dataset. The red dot indicates the observed recurrence rate of the non-psychiatric version of ExAC. (**c**) Allele frequency distribution of class 2 *de novo* CpGs by cohort with the matching distribution of CpG variants in ExAC for comparison. Allele frequency distributions do not significantly differ (*P* = 0.14; Wilcoxon rank sum test). Error bars represent 95% confidence intervals throughout (**a)** – (**c)**. ID/DD, intellectual disability / developmental delay; ASD, autism spectrum disorder

We then sought to determine whether class 1 and class 2 *de novo* variants contribute equally to ASD and ID/DD risk. As a control for the comparison of functional variants, rates of both class 1 and class 2 *de novo* synonymous variants are equivalent across ASD, ID/DD, and unaffected ASD siblings (**Fig. 3A; Supplementary Table 9**) and remain unchanged when we removed the psychiatric cohorts within ExAC (**Supplementary Fig. 3A; Supplementary Table 10**). Thus, collectively neither class 1 nor class 2 *de novo* synonymous variants demonstrated an association with ASD or ID/DD, consistent with previous reports that as a class, *de novo* synonymous variants do not contribute to risk^5-10^. While previous reports implicated *de novo* PTVs as significant risk factors for ASD^5,6,15,16^ and ID/DD^13^, the class 2 *de novo* subset of PTVs show no such association for either ASD (0.015 per case vs. 0.023 per unaffected ASD sibling; *P*=0.98; one-sided Poisson exact test^32^) or ID/DD (0.016 per case vs. 0.023 per unaffected ASD sibling; *P*=0.94; one-sided Poisson exact test), with slightly higher rates in unaffected ASD siblings (**Fig. 3B; Supplementary Table 11**). By contrast, after removing class 2 *de novo* PTVs, class 1 *de novo* PTVs are significantly more enriched in individuals with ASD (0.13 per case) and ID/DD (0.19 per case) as opposed to unaffected ASD siblings (0.07 per control) (ASD vs. control: rate ratio [RR]=1.83; *P*=6.07x10^−12^, ID/DD vs. control: RR=2.61; *P*=6.31x10^−21^; one-sided Poisson exact test). The lack of excess case burden in class 2 *de novo* variants was consistent with what would be expected if such variants did not contribute to ASD and ID/DD risk. However, to ensure we were not losing causal variants by removing all *de novo* variants found in ExAC, we tested the class 2 *de novo* PTVs at three ExAC allele frequency (AF) thresholds: singletons (1 allele in ExAC), AF < 0.0001, and AF < 0.001. We found no significant difference between the rate of class 2 *de novo* PTVs between individuals with ID/DD or ASD as compared to unaffected ASD siblings at any threshold (**Fig. 3C; Supplementary Table 12**). Furthermore, these results remain consistent regardless of whether the psychiatric exomes in ExAC are retained or excluded, demonstrating they are not driving the observed associations (**Supplementary Fig. 3B; Supplementary Table 13**). Thus, the data provides no evidence to suggest these class 2 *de novo* variants contribute to the previously observed enrichment of *de novo* variation in ASD and ID/DD cases, and removing those variants present in ExAC increases the effect size for *de novo* PTVs in ASD and ID/DD. Moving forward, we focus our analyses solely on variation absent from ExAC.

## Gene level analyses

Since observed risk to ASD or ID/DD was carried only by *de novo* variants absent from the standing variation of ExAC, we next sought to extend this concept by evaluating whether the overall rate of PTVs per gene in ExAC provided a similar guide to which ASD and ID/DD variants were relevant. Specifically, we investigated whether the gene-level constraint metric, pLI^16^ (probability of loss-of-function intolerance), could improve our ability to decipher which class 1 *de novo* PTVs confer the most risk to ASD and ID/DD (Online Methods). Using the same threshold as Lek et al. (2016), we used a threshold of pLI ≥0.9 to define loss-of-function (LoF)-intolerant genes and investigated whether individuals with ASD had an increased burden of class 1 *de novo* PTVs in such genes. When we restricted to solely class 1 *de novo* PTVs in LoF-intolerant genes, we observed a significant excess in individuals with ASD (0.067 per exome) compared to their unaffected siblings (0.021 per exome; RR=3.24; *P*=3.14x10^−16^; one-sided Poisson exact test). For individuals with ID/DD, the rate of class 1 *de novo* PTVs in LoF-intolerant genes becomes more striking, with a rate of 0.139 per exome, resulting in a 6.70 RR when compared to the control group of unaffected ASD siblings (P=6.34x10^−38^; one-sided Poisson exact test). By contrast, the rate of class 1 *de novo* PTVs in LoF-tolerant genes (pLI <0.9) show no difference between individuals with ASD (0.063 vs. 0.051; P=0.06; two-sided Poisson test), or individuals with ID/DD (0.048 vs. 0.051; *P*=0.75; two-sided Poisson exact test; **Fig. 3D; Supplementary Table 14**) when compared to unaffected ASD siblings. Again, results remain unchanged when we exclude the ExAC psychiatric samples (**Supplementary Fig. 3C; Supplementary Table 15**). The same trend is observed in congenital heart disease^26,27^ and schizophrenia^25^ (**Supplementary Note; Supplementary Tables 16–21**). Hence, all detectable *de novo* PTV signal in these phenotypes can be localized to 18% of genes with clear intolerance to PTVs in ExAC, with, consequently, substantially amplified rate ratios in this gene set.

**Figure 3.**
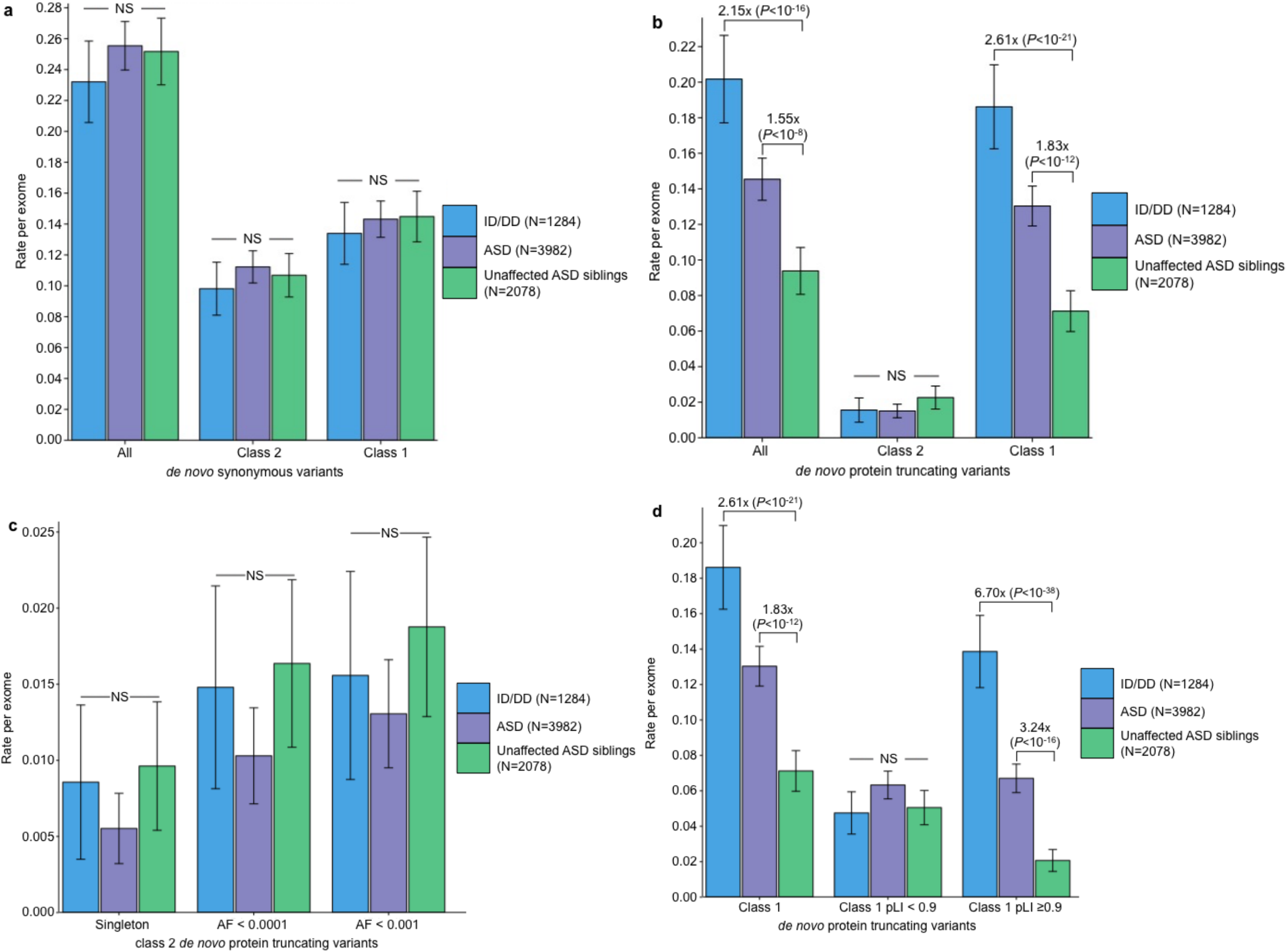
Partitioning the rate of *de novo* variants per exome by class 1, class 2, and pLI. Within each grouping, the rate – variants per individual – is shown for ID/DD (left), ASD (middle), and unaffected ASD siblings (right) with the number of individuals labeled in the legends. (**a**) Rate of *de novo* synonymous variants per exome partitioned into class 2 (middle) and class 1 (right). No significant difference was observed for any grouping of *de novo* synonymous variants. (**b**) Rate of *de novo* PTVs per exome partitioned into class 2 (middle) and class 1 (right). Only class 1 *de novo* PTVs in ID/DD and ASD show association when compared to unaffected ASD siblings. (**c**) Rate of class 2 *de novo* PTVs broken by different ExAC global allele frequency (AF) thresholds: singleton (observed once; left), AF < 0.0001 (middle), and AF < 0.001 (right). (**d**) Rate of class 1 *de novo* PTVs partitioned into class 1 *de novo* PTVs in pLI ≥ 0.9 genes (right), and class 1 *de novo* PTVs in pLI < 0.9 genes (middle). The entire observed association for *de novo* PTVs resides in class 1 *de novo* PTVs in pLI ≥ 0.9 genes. For all such analyses, the rate ratio and significance were calculated by comparing the rate for ID/DD and ASD to the rate in unaffected ASD siblings using a two-sided Poisson exact test^32^ for synonymous variants and one-sided for the remainder (Online Methods). Error bars represent 95% confidence intervals throughout (**a)** – (**d**). See **Supplementary Fig. 3** for the corresponding figures using the non-psychiatric version of ExAC. ID/DD, intellectual disability / developmental delay; ASD, autism spectrum disorder; PTV, protein truncating variant; pLI, probability of loss-of-function intolerance; NS, not significant

Recent studies inferred the presence of multiple *de novo* PTVs in the same gene as evidence of contribution to ASD risk^5-10^. Of the 51 genes with ≥2 *de novo* PTVs, only 38 are absent in controls (**Supplementary Table 22**). This not only reinforces the point that the mere observation of multiple *de novo* PTVs in a gene is not sufficient to define that gene as important^5,16^, but also provides an opportunity to explore whether the pLI metric can refine the identification of specific genes. In fact, 32 of the 38 case-only genes, but only 5 of 13 control-only or case-control hit genes, are LoF-intolerant, a highly significant difference (OR=8.07; *P*=0.003; Fisher’s exact test) that greatly refines the list of genes to be pursued as likely ASD contributors.

## Phenotypic associations for class 1 *de novo* PTVs in LoF-intolerant genes

While enrichment of *de novo* PTVs is one of the hallmarks of ASD *de novo* studies^5-10,15,16^, another consistent finding is an increased burden of these variants among females with ASD^6,15^ and in ASD individuals with low full-scale IQ (FSIQ)^6,15,16^. We investigated whether these hallmarks were present in the 6.55% of ASD cases with a class 1 *de novo* PTV in LoF-intolerant genes (pLI ≥0.9). Indeed, females are overrepresented in the subset (12.26% of females; 5.80% males; *P*=1.75x10^−5^; Fisher’s exact test; Supplementary Table 23). Importantly, for the 6.86% of ASD cases with a class 2 *de novo* PTV or a class 1 *de novo* PTV in a LoF-tolerant gene (pLI <0.9), there is no difference between the sexes, with 6.86% of females and 6.83% of males falling in this category (*P*=1; Fisher’s exact test; Supplementary Table 24). Furthermore, class 2 *de novo* PTVs and class 1 *de novo* PTVs in LoF-tolerant genes show no association with FSIQ (β=-0.001; *P*=0.76; Poisson regression), while class 1 *de novo* PTVs in LoF-intolerant genes predominately explain the skewing towards lower FSIQ (β=-0.023; P=7x10^−8^; Poisson regression; Fig. 4A). Given these observations, we split the ASD class 1 *de novo* PTV signal in LoF-intolerant genes by sex and intellectual disability status (Online Methods). Females with comorbid ASD and intellectual disability have the highest rate of class 1 *de novo* PTVs in LoF-intolerant genes (RR=8.71; P=2.73x10^−12^; one-sided Poisson exact test). Despite the overwhelming enrichment in females and individuals with comorbid ASD and intellectual disability, males with ASD without intellectual disability still show enrichment of class 1 *de novo* PTVs in LoF-intolerant genes (RR=2.95; *P=* 1.31x10^−9^; one-sided Poisson exact test; **Fig. 4B; Supplementary Table 25**). These secondary analyses strongly support the implication of the primary analysis: that collectively, class 2 *de novo* PTVs and class 1 *de novo* PTVs in LoF-tolerant genes have little to no association to ASD or ID/DD and no observable phenotypic impact on those cases carrying them. By contrast, the class 1 *de novo* variants occurring in LoF-intolerant genes contain the association signal and phenotypic skewing observed to date.

**Figure 4.**
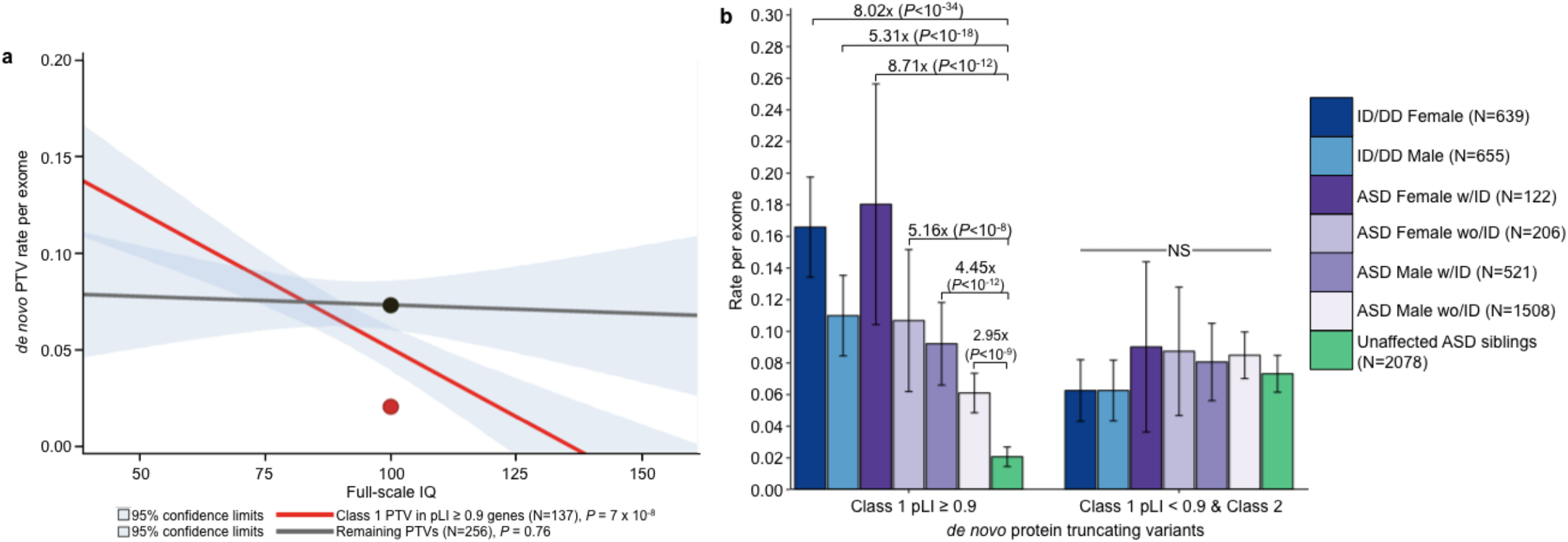
Phenotypic associations for ASD *de novo* PTVs. **(a)** IQ distribution of class 1 *de novo* PTVs in pLI ≥0.9 genes (red) and remaining *de novo* PTVs (class 2 and class 1 pLI < 0.9; grey) in 393 individuals with ASD with a measured full-scale IQ. Dots indicate the rate in unaffected ASD siblings for their respective categories of *de novo* PTVs. P-values calculated using Poisson regression. Only class 1 *de novo* PTVs show association with full-scale IQ. (**b**) Rate of class 1 *de novo* PTVs (left set) and the remaining *de novo* PTVs (class 2 & class 1 in LoF-tolerant genes, right set) in ID/DD (left two bars) and ASD (middle four bars) split by sex and ID with the number of individuals labeled in the legends. Error bars represent 95% confidence intervals, and P-values were calculated using one-side Poisson exact tests comparing to unaffected ASD siblings. ASD, autism spectrum disorder; PTV, protein truncating variant; pLI, probability of loss-of-function intolerance

## Inherited variation

As the effect size for *de novo* PTVs increased after removing those variants present in ExAC, we postulated a similar increase could be obtained from rare inherited PTVs. Under the assumption that risk-conferring variants should be transmitted more often to individuals with ASD, we tested for transmission disequilibrium in a cohort of 4319 trios with an affected proband (Online Methods). Without filtering by pLI or presence/absence status in ExAC, singleton PTVs, as a class, showed no over-transmission (*P*=0.31; two-sided binomial test). After removing all of the variants present in ExAC or in a LoF-tolerant gene (pLI <0.9), we found a modest excess of transmitted PTVs in ASD cases (RR=1.16; *P*=9.85x10^−3^; two-sided binomial test; **Supplementary Table 26**). As with all previous analyses, this result is virtually identical when the psychiatric cohorts in ExAC are removed (RR=1.14; *P*=0.02; two-sided binomial test). While there are far more inherited PTVs than *de novo* PTVs, the inherited variant effect size (1.16 RR) is paradoxically minute by comparison to that of *de novo* PTVs (3.24 RR).

Despite the different effect sizes between *de novo* and inherited PTVs, the data does not suggest the two classes of variation differ in penetrance. Instead, the data suggest the excess of inherited PTVs resides in a different set of genes than those implicated by *de novo* variation. Specifically, the largest *de novo* variant excess resides in a limited and extremely penetrant set of genes that do not contribute substantially to inherited PTV counts. If we consider the 11 genes with ≥3 class 1 *de novo* PTVs in ASD cases and none in controls (47 *de novo* PTVs in total), all 11 are intolerant of truncating variation (pLI ≥0.9) (**Table 1; Supplementary Table 22**). These variants confer risk to particularly severe outcomes: of the cases with available IQ data, 14 of the 29 individuals have IQ below 70 or were unable to complete a traditional IQ test^15^, while only 27% of all ASD individuals with available IQ data in this study fall into this group (*P*=0.008; Fisher’s exact test). Across this same gene set, there are only 4 inherited PTVs (from a total of 5 observed in the parents of the 4,319 ASD trios). Of those 4 inherited PTVs, only the inherited frameshift in *CHD8* bore evidence of mosaic transmission (*P*=5.49x10^−3^; two-sided binomial test; **Supplementary Table 27**) suggesting it may have arisen post-zygotically and not carried by a parent. This ratio – that 80-90% of the observed variants are *de novo* rather than inherited in ASD cases – indicates enormous selective pressure against mutations in these genes, far greater than the direct selection against ASD in general (**Table 1**). Indeed, despite ascertaining these 11 genes based on those with the most class 1 *de novo* PTVs in ASD, we observe a higher rate of *de novo* PTVs in these same genes in the ID/DD studies (37 mutations in 1284 cases). This underscores that selection against these variants likely arises from more severe and widespread impact on neurodevelopment and cognition. Despite the minor contribution of inherited variation in these genes, some insights from studying families may be particularly useful. To our surprise, 1 of the 4 inherited PTVs, a nonsense variant in *ANK2*, was also observed *de novo* in an unrelated individual with ASD, providing a rare instance in which the same variant was observed both inherited and *de novo* in two unrelated individuals with ASD, yet absent from 60,706 individuals in ExAC (**Supplementary Note**).

**Table 1.** Top 12 genes with ≥ 3 class 1 *de novo* PTVs in individuals with ASD.

| Gene | Class 1 *de novo* PTVs |  |  | Inherited |  | Case-control |  | pLI | *P*-value |
| --- | --- | --- | --- | --- | --- | --- | --- | --- | --- |
|  | ASD | Unaffected<br>ASD siblings | ID/DD | T | U | Case | Control |  |  |
| *CHD8* | 7 | 0 | 0 | 1 | 0 | 0 | 0 | 1 | 3.70E-13 |
| *ARID1B* | 5 | 0 | 11 | 0 | 0 | 0 | 0 | 1 | 1.07E-08 |
| *DYRK1A* | 5 | 0 | 2 | 0 | 0 | 0 | 0 | 0.9996 | 2.46E-11 |
| *SYNGAP1* | 5 | 0 | 9 | 0 | 0 | 0 | 0 | 1 | 2.47E-10 |
| *ADNP* | 4 | 0 | 4 | 0 | 0 | 1 | 0 | 0.9989 | 3.93E-09 |
| *ANK2* | 4 | 0 | 0 | 1 | 1 | 0 | 0 | 1 | 7.07E-06 |
| *DSCAM* | 4 | 0 | 0 | 2 | 0 | 1 | 0 | 1 | 3.62E-07 |
| *SCN2A* | 4 | 0 | 7 | 0 | 0 | 1 | 0 | 1 | 1.25E-06 |
| *ASH1L* | 3 | 0 | 0 | 0 | 0 | 0 | 0 | 1 | 1.67E-04 |
| *CHD2* | 3 | 0 | 2 | 0 | 0 | 0 | 0 | 1 | 7.81E-05 |
| *KDM5B* | 3 | 2 | 0 | 5 | 1 | 0 | 0 | 5.09E-05 | 7.22E-05 |
| *POGZ* | 3 | 0 | 2 | 0 | 0 | 2 | 0 | 1 | 3.12E-05 |

12 genes with $\geq 3$ class 1 *de novo* PTVs in 3982 individuals with ASD. Additionally, for each gene, we have listed the number of class 1 *de novo* PTVs in 2078 unaffected ASD siblings and in 1284 individuals with ID/DD, as well as the number of singleton, LOFTEE (see URLs) high-confidence PTVs absent from ExAC that were transmitted (T) or untransmitted (U) to 4319 individuals with ASD and present in 404 cases of ASD and 3654 population controls. *P*-values represent the Poisson probability of observing more than the expected number of class 1 *de novo* PTVs (Online Methods). ID/DD, intellectual disability / developmental delay; ASD, autism spectrum disorder; PTV, protein truncating variant; pLI, probability of loss-of-function intolerance

## Case-control analysis

Having observed a significant enrichment in both *de novo* and inherited PTVs absent from ExAC in LoF-intolerant genes (pLI ≥0.9), we applied this same methodology to case-control cohorts. Given that the variation present in a single individual will be a combination of *de novo* (both somatic and germline) and inherited variation, we expect to see an effect size for PTVs intermediate between that of the *de novo* and inherited PTVs absent from ExAC in LoF-intolerant genes. Using a published cohort of 404 ASD cases and 3654 controls from Sweden^5^, we first analyzed the rate of singleton synonymous variants as a control for further analyses. We found no case-control difference among those present/absent from ExAC (*P*=0.59; Fisher’s exact test; **Supplementary Table 28**). Turning to the PTV category, we observe a slight excess of singleton PTVs in cases with ASD (917 PTVs in 404 cases) compared to controls (7259 PTVs in 3654 controls; OR=1.16; *P*=3.13x10^−5^; Fisher’s exact test; **Supplementary Table 29**). This signal increases once we remove all singleton PTVs present in ExAC or in LoF-tolerant genes, providing the first instance of an exome-wide excess of PTVs demonstrated in ASD without the use of trios (128 PTVs in 404 cases, 447 PTVs in 3654 controls; 2.63 OR; P=1.37x10^−18^; Fisher’s exact test; **Supplementary Table 30 & 31**). Consistent with the previous *de novo* and inherited analyses, no signal exists for the remaining 7601 singleton PTVs (OR=1.06; *P*=0.11; Fisher’s exact test; **Supplementary Table 32**). Lastly, removing the psychiatric cohorts from ExAC results in a 2.42 OR for singleton PTVs absent from ExAC in LoF-intolerant genes (133 PTVs in 404 cases, 506 PTVs in 3654 controls; *P*=1.06x10^−16^; Fisher’s exact test; Supplementary Table 33).

## Discussion

Here we demonstrated that ~1/3 of *de novo* variants identified in neurodevelopmental disease cohorts are also present as standing variation in ExAC, indicating the presence of widespread mutational recurrence. Reinforcing this, we demonstrated that these class 2 *de novo* variants are enriched for more mutable CpG sites. Most importantly, however, these class 2 *de novo* variants confer no detectable risk to ID/DD and ASDs, and eliminating them from our analysis improved all genetic and phenotypic associations by removing the “noise” of benign variation.

We further refined the class 1 *de novo* PTV association using a gene-level intolerance metric (pLI) developed using the ExAC resource and identified that all detectable mutational excess resided in 18% of genes most strongly and recognizably intolerant of truncating mutation. Specifically, 13.5% (±2.0%, 95% CI) of individuals with ID/DD and 6.55% (±0.8%, 95% CI) of individuals with ASD, but only 2.1% (±0.6%, 95% CI) of controls, have a *de novo* PTV absent from ExAC and present in a gene with a very low burden of PTVs in ExAC (pLI ≥0.9). ASD cases with such a variant are more likely to be female and/or have intellectual disability than the overall ASD population. For the remaining 93.45% of the ASD cohort, we fail to observe any meaningful phenotypic difference (i.e., IQ or sex) between the 6.86% of individuals with and the 86.59% of individuals without a class 2 *de novo* PTV or a class 1 *de novo* PTV in a LoF-tolerant gene. These results, taken together with an overall lack of excess case burden, suggest that collectively, neither class 2 nor class 1 *de novo* PTVs in LoF-tolerant genes (pLI <0.9) appear to confer significant risk toward ASD. Thus, we have refined the role of *de novo* protein truncating variation in ASD, confining the signal to a smaller subset of patients than previously described^6,33^.

This analysis framework, operating at the variant level, also enabled a careful examination of inherited variation in ASD. While ASD is highly heritable^3^, few analyses^34^ have demonstrated specific inherited components. By removing inherited PTVs present in ExAC or in LoF-tolerant genes, we discovered a modest signal of over-transmitted PTVs, in line with previous reports^34^. The vast majority of inherited PTVs appear to affect genes that have yet to show signal from *de novo* variation, with only 1% residing in the strongest associated genes, indicating the inherited variants reside in genes with a somewhat weaker selective pressure against them. Ultimately, however, as these variants occur in 15.4% of cases but carry only a 1.16-fold increased risk as a group, they explain little of the overall heritability (<1% of the variance in liability).

Given the current size of ExAC and the general scarcity of truncating variants, the pLI metric for constraint against loss-of-function variation does not yet provide precise resolution of the selection coefficient acting on PTVs in that gene. That is, even a pLI ≥0.9 does not guarantee a selection coefficient sufficiently high to ensure the vast majority of variation is *de novo* rather than inherited. In fact, selection coefficients for pLI ≥0.9 genes range from 0.1–0.5 (where the majority of variation will be inherited), all the way to selection coefficients approaching 1, in which the variants are almost completely reproductively null. Only larger reference panels will enable refining these estimates, articulating a gradient from the strongest genes we currently flag (e.g., the 11 genes with ≥3 *de novo* PTVs in ASD and none in controls that make their contribution almost entirely through penetrant, single-generation *de novo* variation) to those genes we have yet to define clearly that will make their contribution largely through inherited, albeit less penetrant, variation. The significant expansion of exome sequencing in ASD, alongside larger reference panels from which to draw more precise inferences about selective pressure against variation in each gene, will allow us to fill in the genetic architecture of ASDs in the region of the effect size spectrum between severe *de novo* variation at one end and common variation at the other.

ExAC currently has 15,330 individuals from psychiatric cohorts, with the schizophrenia cohort being the largest24. Given the shared genetics between ASD and schizophrenia ^2,5,16,17,25,35,36^, it is reasonable to hypothesize that the psychiatric cohorts within ExAC could influence our analyses. As we have shown however, removing the psychiatric cohorts within ExAC does not change our results. In fact, of the 16 *de novo* PTVs in LoF-intolerant genes that were also variant in ExAC, only two reside solely in the 15,330 individuals from the psychiatric cohorts *(CUX2* in ASD, *LARP1* in unaffected ASD siblings). This number being so small is in retrospect not surprising because it is so unusual to observe a deleterious variant both *de novo* and present as standing variation in individuals with the same ascertained phenotype, let alone in different ascertained phenotypes. The *ANK2* nonsense variant was the only such instance of the same deleterious variant being *de novo* in one ASD trio and inherited in another.

While we use ASDs and ID/DD here to explore this framework, it can certainly be applied toward any trait. However, this framework is optimally powered in traits governed by genes under strong selection, as it will remove *de novo* variants that are more common when examined in the context of a larger reference population. Our results reinforce the point that not all *de novo* variants are rare and contribute to risk, while highlighting the tremendous value of large population sequence resources even for the interpretation of *de novo* variation and complex disease. This is especially important in the case of clinical sequencing, in which the paradigm has unfortunately become that if a protein-altering *de novo* variant is present in the gene of interest, then it is often considered the causal variant^37,38^. Clearly, not all *de novo* variants are equal, and not all *de novo* variants in a gene contribute to risk in the same way.

## Online Methods

### Datasets and data processing

Two versions of the Exome Aggregation Consortium (ExAC) database were used in this analysis: the full version of ExAC (N = 60,706) and the non-psychiatric version of ExAC (N = 45,376). The non-psychiatric version of ExAC has the following cohorts removed: Bulgarian trios (N = 461), sequencing in Suomi (N = 948), Swedish schizophrenia & bipolar studies (N = 12,119), schizophrenia trios from Taiwan (N = 1505), and Tourette syndrome association international consortium for genomics (N = 297). We used a combined set of 8401 published *de novo* variants from 3982 probands with ASD and 2078 of their unaffected siblings from two recent large-scale exome sequencing studies: de Rubeis *et al* (N_ASD_ = 1474, N_una_ffecte_d sib_ = 267)^5^, Iossifov, O’Roak, Sanders, Ronemus *et al* (N_ASD_ = 2508, N_unaffected sib_ = 1911)^6^ (**Supplementary Table 1**). We also used 1692 *de novo* variants from 1284 probands published in studies of intellectual disability (ID) (de Ligt *et al:* N = 100^12^, Rauch *et al:* N = 51^14^) and developmental delay (DD) (DDD: N = 1133)^39^ (**Supplementary Table 2**). *De novo* variants from congenital heart disease^26,27^ and schizophrenia_25_ were also downloaded for additional confirmation of the recurrent mutation rate (**Supplementary Tables 5 and 6**). Details of the sequencing and *de novo* calling can be found in the referenced publications.

To ensure uniformity in variant representation and annotation across datasets and with respect to the ExAC reference database^40^, we created a standardized variant representation through a Python implementation of vt normalize^41^ and re-annotated all variants using Variant Effect Predictor (VEP)^42^ version 81 with GENCODE v19 on GRCh37. VEP provided the Ensembl Gene IDs, gene symbol, the Ensembl Transcript ID for use in determining canonical transcripts, as well as PolyPhen2 and SIFT scores. We used the canonical transcript when possible for cases when the variant resided in multiple transcripts, and the most deleterious annotation in cases of multiple canonical transcripts. If no canonical transcript was available, the most deleterious annotation was used. As such, variants in Supplemental Tables 1, 2, and 6 may differ from their respective publications due to standardizing variant representation and annotation.

## Determining class 1 or class 2 *de novo* variants

*De novo* variants were classified as class 1 or class 2 based on their respective absence or presence in ExAC. Presence or absence in ExAC was defined if the variant had the same chromosome, position, reference, and alternate allele in both files. Due to the heterogeneous nature of ExAC, and the different capture arrays used in the original exome sequencing studies incorporated into ExAC, we elected to use all of the variants in ExAC, not just those with a PASS status in the GATK variant calling filter. For insertions/deletions, we took a conservative stance that they must match exactly (i.e., a subset was not sufficient). To illustrate, if a *de novo* variant on chromosome 5 at position 77242526 has a reference allele of AGATG and a *de novo* alternate allele where four nucleotides are deleted (AGATG to A), we would not say that variant is present in ExAC if there was another variant at the same genomic position in ExAC where only the first two of these nucleotides are deleted (AGA to A). Lastly, for variants outside of the proportion of the genome covered by ExAC, we considered them to be class 1 *de novo* variants – as expected, none of these variants reside in the coding region (**Supplementary Table 34**).

## Variant calling for transmission and case-control analysis

We used the Genome Analysis Toolkit (GATK v3.1-144) to recall a dataset of 22,144 exomes from the Autism Sequencing Consortium (ASC)^43^ & Simons Simplex Collection (SSC) ^44^ sequencing efforts. This call set contained 4319 complete trios (including all those from which the published and validated *de novo* mutations were identified), which we used to evaluate inherited variation, and a published case-control dataset of individuals of Swedish ancestry (404 individuals with ASD and 3564 controls)^5^. We applied a series of quality control filters on the genotype data, using the genome-wide transmission rate as a guide for filter inclusion/exclusion. More specifically, we calibrated various genotyping filters such that synonymous singleton variants – where the alternative allele was seen in only one parent in the dataset – was transmitted at a rate of 50%, because we expect, as a class, synonymous variants to be transmitted 50% of the time. As with the ExAC analysis^40^, we found GATK’s default Variant Quality Score Recalibration (VQSR) too restrictive due to the bias toward common sites. In order to reduce the number of singleton variants being filtered out, we recalibrated the Variant Quality Score Log Odds (VQSLOD) threshold from -1.49 to -1.754, dropping the singleton synonymous transmission rate from 51.1% to 49.9998% (**Supplementary Fig. 4**). Additional filtering was done at the individual level, in which we required a minimum read depth of 10 and a minimum GQ and PL of 25 for each individual’s variant call. We also applied an allele balance filter specific for each of the three genotypes (homozygous reference, heterozygous, homozygous alternate), where allele balance is defined as the number of alternate reads divided by the total number of reads. We required the allele balance for homozygous reference individuals to be less than 0.1, allele balance for heterozygous individuals to be between 0.3 and 0.7, and the allele balance for homozygous alternate individuals to be greater than 0.9. Calls that did not pass these filters were set to missing. Lastly, for the transmission analysis, we removed variants in which more than 20% of families failed one of our filters. For the case-control analysis, we removed variants in which more than 5% of families failed one of our filters.

## On the use of the Poisson exact test for comparing rates of *de novo* variation between two samples

As with many other papers^6,8,45-47^, we too were interested in testing whether the rate of a given class of *de novo* variation was significantly different between our cohorts of individuals with ASD or ID/DD as compared to unaffected ASD siblings. As the number of *de novo* variants per individual follows a Poisson distribution^8^, we tested *H_A_* : *λ*_1_ ≠ *λ*_2_ vs. *H*_0_ : *λ*_1_ = *λ*_2_, where *λ_i_* is the rate of a given class of *de novo* variation in group i, using the Poisson exact test (also known as the *C*-test)^32^. Note: we could not compare the rates to expectation, because the expectations published in Samocha et al., (2014) are for ALL *de novo* variants, not just *de novo* variants present/absent from ExAC. An important consequence of our hypothesis test is that effect sizes are reported as rate ratios (RR), which is simply the quotient of the two rates. While more commonly reported, odds ratios require Bernoulli random variables (e.g., an individual either harbors or does not harbor a *de novo* variant), and as such, would be incorrect given the hypothesis we are testing. Had we been interested in testing for a significant difference between the proportion of individuals harboring a *de novo* PTV, then an odds ratio would be appropriate (and Fisher’s exact test would suffice in this case). Thus, only in using the Poisson exact test could we reject the null hypothesis that the rate of *de novo* PTVs is the same between individuals with ASD and their unaffected siblings and find evidence that individuals with ASD have a higher rate of *de novo* PTVs than their unaffected siblings. The difference between the two tests is a subtle, but important one.

## On the use of pLI (probability of loss-of-function intolerance)

Using the observed and expected number of PTVs per gene in the ExAC dataset, we developed a metric to evaluate a gene’s apparent intolerance to such variation^24^. Briefly, the probability of loss-of-function intolerance (pLI) was computed using an EM algorithm that assigned genes to one of three categories: fully tolerant (in which PTVs are presumed neutral and occur, like synonymous variants, at rates proportional to the mutation rate), “recessive-like” (showing PTV depletion similar to known severe autosomal recessive diseases) and “haploinsufficient-like” (showing PTV depletion similar to established severe haploinsufficiencies). pLI is the posterior probability that a gene resides in the last, most loss-of-function intolerant, category. See section 4 of the supplement of Lek, et al. (2016) for more details.

## Phenotype Analysis

Full-scale deviation IQ scores were measured using several tests including the Differential Ability Scales, the Wechsler Intelligence Scale for Children, and the Wechsler Abbreviated Scale of Intelligence. IQ has previously been associated with *de novo* PTV rate in the SSC^6,15,48^. In this analysis, we used Poisson regression to estimate the relationship between the rate of each of class 1 and class 2 PTVs and proband full-scale deviation IQ.

## Calculating the expected number of class 2 *de novo* variants in a reference database

For a set of r *de novo* variants, each with the same allele count, K, in ExAC, we can estimate the number of those variants still observed at least once in a subset of size n using the hypergeometric distribution. That is to say, how many of those same sites will still be present as standing variation in a down-sampled version of ExAC? Specifically,

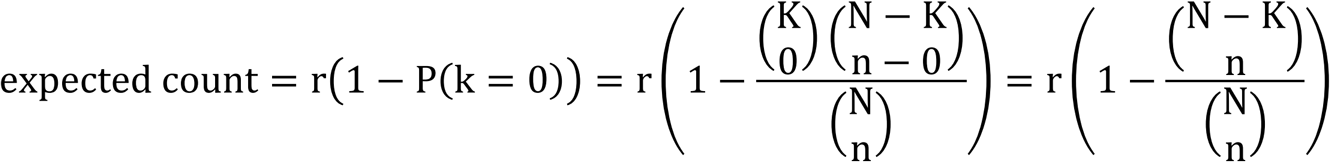

where k is approximatelyhypergeometric (N, K, n), and N is the number of chromosomes in the current version of ExAC (N=121,412).

This only holds when each down-sampled set of ExAC preserves the ancestry proportions of the total sample.

## Calculating mutation rates for class 1 and class 2 *de novo* PTVs

Samocha et al. calculated per gene mutation rates for ALL synonymous, missense, and PTVs, not for those present/absent in ExAC. If we are interested in comparing the rate of class 1 *de novo* PTVs to the expected depth-corrected mutation rate for class 1 *de novo* PTVs, we can roughly calculate it. For a given gene, we can derive the class 1 and class 2 PTV mutation rate by breaking down the overall mutation rate for PTVs, denoted as 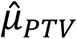, using equation (1)

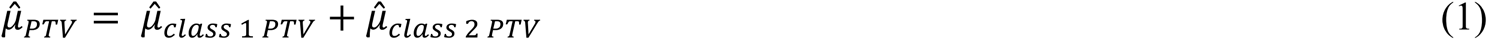

In case the logic behind equation 1 isn’t completely clear, it may help to point out that the number of class 1 and class 2 PTVs is equal to the total number of PTVs. Now, Samocha et al. provides us with 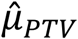, so all we need to do is calculate 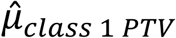 and 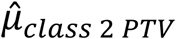. Given all of the PTVs in ExAC, and the probability of each trinucleotide-to-trinucleotide mutation, we can calculate 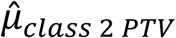 using equation (2)

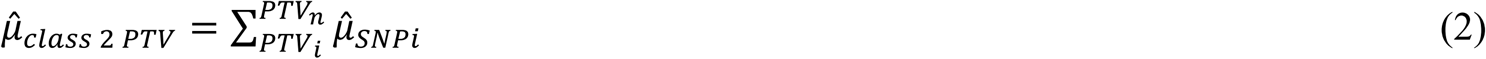

where *i* indexes the *n* PTVs for a given gene present in ExAC, and 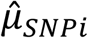 is the mutation rate of that specific trinucleotide substitution that creates a PTV. With 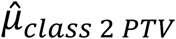 calculated, 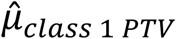 follows from equation 1. However, these per gene 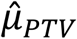 calculations do not account for sequencing depth. Correcting for depth of sequencing becomes tricky, as the depth of sequencing varies between studies and will not necessarily be the same as the depth of sequencing for ExAC. However, we can roughly approximate the depth-corrected 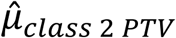 for each gene using the following equation under the assumption that the fraction of the raw mutability from class 2 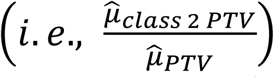 is equal to the fraction of the class 2 depth-corrected mutability 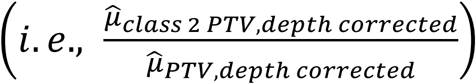

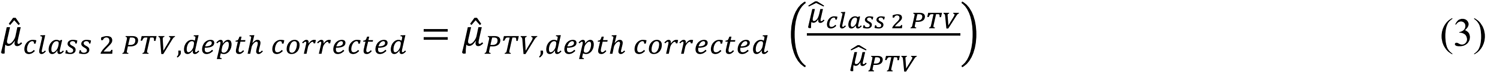

The depth corrected 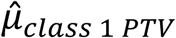 follows using the same logic as we used in equation (1).

## Data availability & accession codes

Data included in this manuscript is deposited in the database of Genotypes and Phenotypes (dbGaP) under accession phs000298.v2.

## URLs

Exome Aggregation Consortium (ExAC), http://exac.broadinstitute.org/; LOFTEE, https://github.com/konradjk/loftee

## Acknowledgements

We would like to thank all of the members of the ATGU and the Wall Lab for their assistance in this endeavor. We thank the families who took part in the Simons Simplex Collection study and the Simons Variation in Individuals Project, as well as the clinicians who collected data at each of the study sites. The authors would like to thank the Exome Aggregation Consortium and the groups that provided exome variant data for comparison. A full list of contributing groups can be found on the ExAC website (see URLs). We would also like to greatly thank A. Byrnes, R. Fine, D. Fronk, A. Martin, C. Nichols, N. Radd, K. Satterstrom, and E. Wigdor for their insightful contributions. Lastly, we would like to acknowledge G. A. Barnard for inspiring us to write in a more conversational tone, similar to his seminal 1947 paper, *Significance Tests For 2x2 Tables.* This work was supported by NIH grants U01MH100233, U01MH100209, U01MH100229 and U01MH100239 to the Autism Sequencing Consortium (ASC), and R56 MH097849 and R01 MH097849 to the Population-based Autism Genetics and Environment Study (PAGES). M.J.D., J.A.K., and K.E.S. were supported by grants from the Simons Foundation Autism Research Initiative (SFARI 342292 and a subaward from the Simons Center for the Social Brain at MIT). ML and DGM’s work on the ExAC project was funded by U54DK105566 and R01 GM104371 from the National Institutes of Health. KS was funded by T32 HG002295/HG/NHGRI. EBR was funded by National Institutes of Mental Health Grant 1K01MH099286 and NARSAD Young Investigator grant 22379.

## Author Contributions

JAK and EBR performed the analyses. JAK, DPH, EBR, and MDJ designed the experiment. JAK and KS wrote the code. DPW, EBR, and MJD supervised the research. JAK and MJD wrote the paper. JAK, KES, DPH, SJS, ML, KJK, DGM, and JDB generated data. JAK was responsible for the remainder. All authors revised and approved the final manuscript.

